# Chronic temperature stress effects on the liver proteome of two threespine stickleback (*Gasterosteus aculeatus*) populations using a novel DIA assay library

**DOI:** 10.1101/2021.06.21.449281

**Authors:** Bryn Bo Levitan, Dietmar Kültz

**Affiliations:** Department of Animal Sciences, University of California Davis, Davis, CA

**Keywords:** proteomics, chronic temperature acclimation, liver, threespine stickleback

## Abstract

A data-independent acquisition (DIA) assay library was generated for the liver of threespine sticklebacks to evaluate alterations in protein abundance and functional enrichment of molecular pathways following either chronic warm (25°C) or cold (7°C) three-week temperature challenge in two estuarine populations. The DIA assay library was created from a data-dependent acquisition (DDA) based raw spectral library that was filtered to remove low quality or ambiguous peptides. Functional enrichment analyses using STRING identified larger networks that were significantly enriched by examining both the entire liver proteome and only significantly elevated or depleted proteins from the various comparisons. These systems level analyses revealed the unique liver proteomic signatures of two populations of threespine sticklebacks acclimated to chronic temperature stress. The Big lagoon population (BL) had a stronger response than the Klamath river population (KL). At 7°C, BL showed alterations in protein homeostasis that likely fueled a higher demand for energy, but both populations successfully acclimated to this temperature. The warm acclimation induced major increases in proteins involved in chromatin structure and transcription, while there were decreases in proteins related to translation and fatty acid metabolism. Functional enrichment analyses of the entire liver proteome uncovered differences in glycolysis and carbohydrate metabolism between the two populations and between the cold acclimated and control groups. We conclude that the synchronous regulatory patterns of many proteins observed in the liver of threespine sticklebacks provide more comprehensive insight into population-specific responses to thermal stress than the use of less specific pre-determined biomarkers.

## INTRODUCTION

Temperature is one of the most important abiotic factors that profoundly affects molecular, cellular, and organismal processes including directly fitness-related traits such as reproduction, development, and survival (1–4). Fish, along with reptiles, amphibians, and invertebrates, are mostly ectothermic, thus temperature exerts more control than any other abiotic factor on internal processes (5). It is projected that global temperatures will continue to rise throughout the 21^st^ century along with the duration, intensity, and spatial extent of heat waves (6). Temperature (thermal stress) is likely to be a driver of natural selection (3) and different species, or different populations within a species, are adapted to their unique environmental conditions such that proteins function best at temperatures that match their habitats (7). Coastal ecosystem biodiversity and ecosystem functioning and services have already been impacted by intensified heatwaves, acidification, sea level rise and changes in oxygen and salinity levels (8). In marine and estuary environments, increases in air and water temperatures change biogeographic patterns (9, 10), and populations from different parts of a species’ biogeographic range handle these temperature changes differently (11). Because external temperatures dictate internal temperatures for ectotherms, behavioral modifications are used to a large extent to control their body temperatures, but such means depend on an accessible temperature gradient. Estuaries are especially susceptible to warming, with lagoons and rivers facing the highest levels of warming due to shallower depths and limited exchange with the ocean, which limits opportunities for behavioral modifications (12). Since most fish do not actively regulate temperature via internal mechanisms, temperature is of vital importance for basic physiological functions, and certain habitats, such as estuaries, present greater challenges for escape or migration. One major question of significance given the predictions about climate change is how organisms living across different environmental conditions will respond to increased environmental challenge. Do different populations employ different mechanisms and pathways to regulate internal conditions?

Many species have been used as model organisms for examining thermal stress responses, however, the threespine stickleback (*Gasterosteus aculeatus*) represents an ideal candidate for numerous reasons. Threespine sticklebacks are widely distributed throughout the northern hemisphere, representing many phenotypically diverse populations along both a longitudinal (North America, Europe, Asia) and latitudinal (Mexico to Alaska) gradient (13). These euryhaline fish inhabit freshwater, brackish water, and coastal marine habitat, including habitats most susceptible to warming from climate change such as lagoons and rivers (12). In addition, the genome sequence and a high-quality annotated reference proteome are available for this species. Furthermore, these fish are abundant, easy to capture, survive captivity well, and have relatively short life cycles.

Proteomics is a powerful tool for examining the effects of environmental conditions on organisms. Proteomics arose in the 1980s and was originally developed to allow for the study of the proteome, which represents all the proteins that are expressed by a genome (14, 15). Data-dependent acquisition (DDA) is a proteomics method for selecting the highest abundance precursor ions in MS1 spectra to fragment for MS2 acquisition, thus producing tandem (MS/MS) mass spectra that are then matched to a database for identification (16, 17). Because the most abundant precursor ions are chosen, this is a stochastic method and there can be differences in spectra matched from one run to another, even on the same sample (18). To overcome this stochasticity, data-independent acquisition (DIA) is a more recent method that fragments all precursor ions within a specified m/z window (17). This method allows for precise and reproducible identification and quantification of peptides, including lower abundance precursor ions, and greatly increases the number of proteins that can be reproducibly quantified (19). Protein abundance, synthesis, degradation, protein-protein interactions, location in subcellular compartments, and post-translational modifications such as phosphorylation, glycosylation, acetylation, and methylation can all be examined using proteomics (20–22). Since natural selection ultimately acts on phenotypes, proteins represent a more direct readout of what is being selected for than either the genome or the transcriptome (23). Additionally the correlation between the abundance of transcripts and the corresponding proteins is often highly nonlinear (23–25). Proteomics studies help elucidate which transcript changes are resulting in changes at the protein level. Furthermore, co-expression patterns of proteins can be used to identify which molecular pathways are up- or down-regulated under a given stressor (22). Proteomic analysis of chronic exposure from the laboratory or the field is still sparse (26), even though proteomic signatures to environmental stress exposures provide deep insight into the evolution of organisms to changing environments (27).

In this study, the liver proteomes of two populations of threespine sticklebacks were compared after exposure to either chronic warm or cold stress to understand the proteins and pathways utilized to overcome temperature stress. The liver provides a good representation of the general condition of a fish as it plays a vital role in critical physiological processes such as homeostasis and metabolism of lipids, glucose, and amino acids, detoxification, and immune system function (28, 29). Warm exposure increases metabolic rates, stimulates physiological and behavioral processes, and requires maintenance to counteract protein denaturation, DNA mutations, oxidative damage and cell death, all of which require more energy to sustain at the expense of growth, reproduction, and immunity (30). Cold exposure decreases metabolic rates, alters lipid homeostasis and metabolism, and increases protein degradation, while oxidative stress, altered protein homeostasis, and large metabolic changes are shared responses to temperature stress regardless of directionality (26, 31). However, there are few proteomics experiments examining chronic acclimation to temperature challenge in fish (26, 27). This study examined differences in the protein abundance and functional enrichments in the liver proteome of threespine stickleback populations from two estuarine habitats (lagoon and river) after a three-week chronic acclimation to either 7°C (cold) or 25°C (warm) to characterize population-specific proteomic signatures of chronic temperature stress.

## MATERIALS AND METHODS

Experimental work was approved by and conducted according to UC Davis Institutional Animal Care and Use Committee (IACUC) rules and regulations (IACUC number 18010, AAALAC number 127 A3433-01).

### Breeding of wild-caught fish and rearing of F1 progeny

Fish were collected from Klamath river (salinity: 0 g/kg; temperature: 13.2 °C) in Klamath, CA, and Big lagoon (salinity: 8.3 g/kg; temperature: 14.8 °C) in Trinidad, CA in the fall of 2016. Fish were fed a rotating diet of frozen blood worms, daphnia, and mysis shrimp (Cobalt Aquatics) with 25% water changes three times a week and 12hr light/12hr dark cycle. Fish were kept at 2-3 g/kg salinity and room temperature (16-18 °C). To induce breeding, water temperature was raised to 20°C using a 50W submersible aquarium heater and the light/dark cycle was changed to 16h light/8h dark cycle. Big lagoon (BL) and Klamath river (KL) wild-caught sticklebacks were externally fertilized and hatched in late winter and early spring of 2017. Hatchlings were kept at 0 g/kg and 18°C. Hatchlings were fed live brine shrimp nauplii (San Francisco Bay Brand or E-Z Egg) once or twice a day *ad libitum*. After hatchlings were large enough to be moved (~3 months), they were transitioned to frozen brine shrimp larvae (Hikari) and progressively to daphnia, mysis shrimp, and blood worms (Cobalt Aquatics). Fish were reared for at least 12 months prior to inclusion in any experiments.

### Chronic acclimation experiments

First generation (F1) sticklebacks from BL (N=30) and KL (N=30) were pre-acclimated for three weeks at 15°C and 9 g/kg (plasma-isosmotic conditions that minimize energy expenditure for osmoregulation) prior to experimentation. Fish were fed *ad libitum* once per day during the pre-acclimation and experimental phases after which excess food and waste was removed. The temperature was either increased or decreased from 15°C by 2°C/day up to 25°C for the chronic warm and down to 7°C for the chronic cold acclimation, after which fish were held for 21 days at the respective temperatures. Ten fish from each population were randomly assigned to each experimental group: 25°C (warm), 15°C (control), 7°C (cold). A water bath method was used with pumps and air stones to circulate heated or cooled water around individually aerated jars containing the fish. Heating to 25°C was achieved using 200-watt electric heaters and cooling to 7°C was done using a refrigerated/heated 6L circulating bath (PolyScience, 9106A11B). Controls were kept at 15°C but handled in the same manner as the 25°C and 7°C groups. Fish were held at the final temperatures (25°C warm; 7°C cold) for a total of three weeks. Because the temperature change was 10°C overall for warm and 8°C overall for cold, the 7°C acclimation group was dissected a day earlier than the 25°C acclimation group. Half of the controls were dissected each day to account for any differences between the two days, and the dissections occurred under the same conditions and at the same time of day. Dissection order was alternated among the different conditions and between the two populations. Wet weight, standard length, and lateral plate count were recorded prior to dissection. Fish were sexed upon dissection. Tissues were extracted and individually flash frozen in liquid nitrogen. There was no mortality during this experiment. Figure 1 depicts a schematic overview of the experimental design.

**Figure 1.**
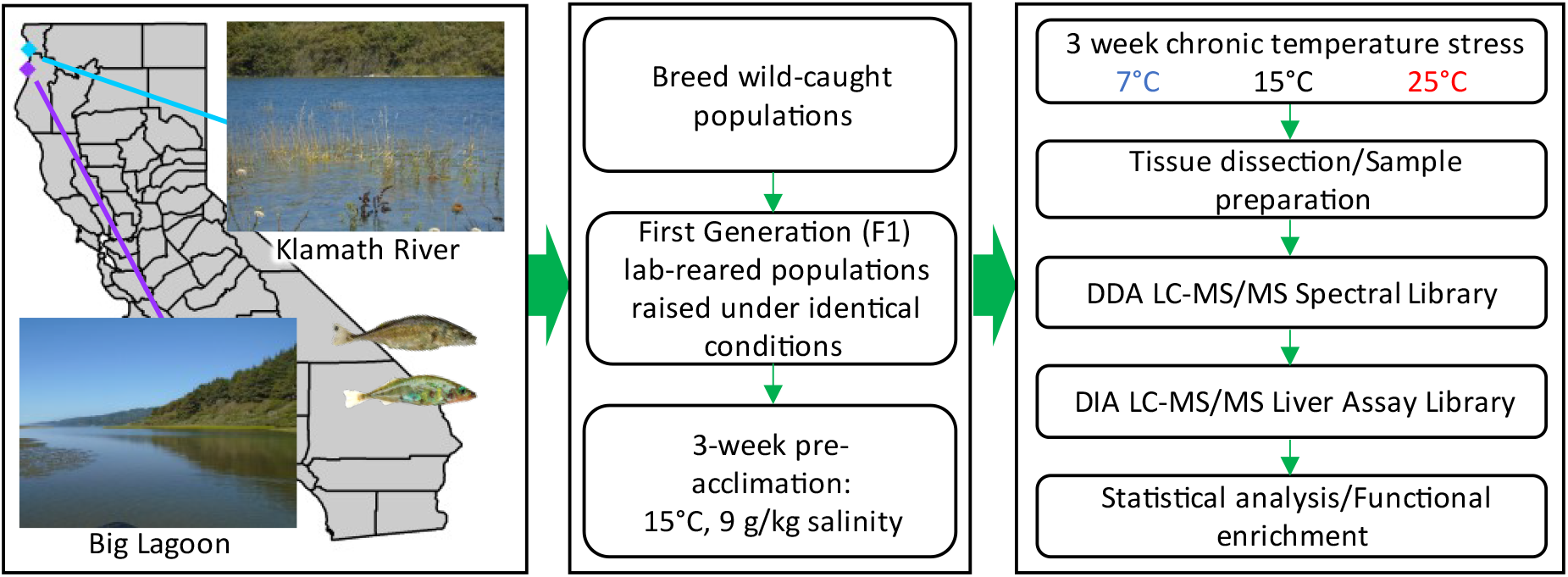
Sampling locations for two wild-caught populations of threespine sticklebacks from Northern California: Klamath river (coordinates: −124.071111, 41.545278) and Big lagoon (coordinates: −124.105994, 41.177013). Wild caught fish were externally fertilized and reared for at least one year under identical conditions in the laboratory, pre-acclimated for three weeks, and then exposed to a three-week chronic temperature stress experiment at either cold (7°C), control (15°C), or warm (25°C) conditions. Liver samples were analyzed by liquid chromatography tandem mass spectrometry (LCMS2) and data transformed into a DIA assay library used to quantify proteins and detect significant abundance differences and functional enrichment.

### Sample preparation

Protein extraction and trypsin digestion were performed as previously documented (32) but with the following modifications detailed below. The liver tissues were crushed using a steel-handle Teflon pestle inside of a 2 mL low retention microcentrifuge tube (LR-MCF) that was dipped in liquid nitrogen. Proteins were reduced with a 150mM dithiothreitol (DTT) stock solution to achieve a final concentration of 10 mM and incubated at 37°C for 30 minutes. Proteins were alkylated with a 450 mM iodoacetamide (IAA) stock solution to a final concentration of 30 mM and incubated in the dark at room temperature for 30 minutes. Proteins were precipitated with an ice-cold solution of 10% trichloroacetic acid (TCA), 0.15% DTT, 90% acetone (5x volume), incubated at −30°C for 30 minutes to precipitate proteins and dissolve contaminants and then centrifuged for 5 minutes at 19,000 g at 4°C. The precipitated pellet was then washed once with an ice-cold solution of 0.15% (weight/volume) DTT in acetone and centrifuged for 5 minutes at 19,000 g at 4°C. The protein pellet was dissolved in a solution of UT buffer (7 M urea/2 M thiourea/0.2% DTT, 6x the volume of the original tissue weight), incubated at room temperature on a rotator for 30 minutes to maximize protein dissolving, centrifuged for 5 minutes at 19,000 g, and the supernatant was removed and transferred to a clean 0.5 mL LR-MCF tube. A 660 nm protein assay (Thermo-Pierce, cat. 22660) compatible with diluted UT buffer was completed in duplicate for each sample. Based on the average protein concentration, 1M ammonium bicarbonate (Ambic) buffer (pH 8.5) and LCMS water were added to dilute samples to 150 ng/100 μL total protein concentration and 100 mM Ambic. Immobilized trypsin beads (Promega, cat. V9012) were added at a 1:30 ratio relative to the total protein and samples were incubated at 35°C for 16 hours on a rotator. The immobilized trypsin was pelleted via centrifugation at 19,000 g for five minutes at 4°C and the supernatant was carefully removed and transferred to a clean LR-MCF tube. Samples were dried by speed vacuum (Thermo-Savant, ISS-110) until just dry, resuspended in 0.1% formic acid (FA), transferred to maximum recovery glass vials (Waters, cat. 186000384C), and stored at 4°C prior to sample injection in the mass spectrometer.

Tryptic peptides (2 μl, 150 ng/μl) were trapped for 1 minute at 15 μL/min on a Symmetry trap column (Waters, cat. 186003514) and separated on a 1.7 μm particle size BEH C18 column (250mm x 75μm, Waters, cat. 186003545) after injection using a nanoAcquity sample manager (Waters, Milford, MA) via reversed-phase liquid chromatography with a nanoAcquity binary solvent manager (Waters). Peptide elution occurred over a linear acetonitrile gradient (3-35%) for 140 minutes directly into a UHR-qTOF mass spectrometer (Impact II, Bruker) using a pico-emitter tip (New Objective FS360–20-10-D-20, Woburn, MA). Samples were run in batches using Hystar 4.1 (Bruker). A 68 fmol BSA peptide mix was run a minimum of once per week to serve as a control to monitor baseline instrument performance.

### Data-dependent acquisition (DDA)

Peak lists were generated using DataAnalysis 4.4 (Bruker Daltonics) from DDA raw data and imported into PEAKS X (Bioinformatics Solutions Inc., Waterloo, Canada). Peptide spectrum matches were identified using PEAKS X and X!Tandem Alanine (The Global Proteome Machine Organization). Unambiguous assignment of peptides to unique proteins was made using the *G. aculeatus* proteome database downloaded from UniprotKB on July 14, 2019. The proteome database included 27,249 *G. aculeatus* proteins plus the same number of randomly scrambled decoys and 282 common contaminants (human keratins, porcine trypsin, etc.). Cleavage specificity for trypsin was C-terminus of either Lys or Arg except when followed by Pro, and up to two missed cleavages were allowed. In the first search round, the following PTMs were allowed: Cys carbamido-methylation, Met oxidation, N-terminal acetylation, and Pro oxidation. Second round PEAKS-PTM searches included all 313 variable PTMs contained in the PEAKS database, with a maximum of two PTMs per peptide allowed. Precursor mass tolerance limits were 20 ppm and fragment ion mass tolerance limits were 0.03 Da. All DDA data are available at MassIVE (MSV000087672) and ProteomeXchange (PXD026823).

### Construction of raw spectral library and DIA assay library

Peptide-to-spectrum matches (PSMs) and protein annotations from the DDA data were exported from PEAKS X in pepxml format and compiled into a raw liver spectral library for *G. aculeatus* in Skyline 20.0 (33). The target list of proteins went through multiple filtering steps as detailed in the first results section (see below). The final DIA assay library and all relevant metadata are available at Panorama Public (https://panoramaweb.org/bbl02.url).

### Data-independent acquisition (DIA)

All samples were run a second time to acquire DIA data. Liquid chromatography conditions were identical to those used for DDA data acquisition, but only MS2 spectra were acquired. The mass range was 390-1015 m/z with a scan rate of 25 Hz in 2.5 second intervals and an isolation width of 10 m/z (1 m/z overlap) as previously described (19).

### Statistical analysis and data visualization

Heat maps were generated using Genesis 1.8.1 (34). Functional enrichment networks were analyzed and created in STRING 11.0 (https://string-db.org) (35). STRING settings were set as follows: Network edges were set to confidence (line thickness indicates strength of data support), all active interaction sources were included (text mining, experiments, databases, co-expression, neighborhood, gene fusion, and co-occurrence), the minimum required interaction score was medium confidence (0.400). Volcano, mass error, retention time, and q-value plots were generated in Skyline 20.1.0.76 (33). Skyline 20.0 was used for quantitative analyses and visualization of DIA data, and slight variations in retention time across runs were corrected using 14 manually selected iRT standards (33). mProphet was used in Skyline to train models that optimized selection of correct peaks (33, 36). The mass accuracy was set at 20 ppm for transitions. For group comparisons, the normalization method employed was equalize medians, the confidence level was 95% at the protein level, the summary method was Tukey’s median polish, and the q-value cutoff was 0.05.

### Functional enrichment analysis

Functional enrichment analysis was conducted with STRING 11.0 (35). For the four overall comparisons (KL vs. BL, 15°C vs. 7°C, 15°C vs. 25°C, and 7°C vs. 25°C) the STRING “proteins with values/ranks” search function was used, with fold change serving as the rank used for the search. This list included the entire protein set with the corresponding fold changes based on the particular comparison for the liver tissue after both automated and manual curation of the assay library. For smaller comparisons (all others), the STRING “multiple proteins” search function was used for significantly up- or down-regulated groups of proteins from that comparison. For all comparisons functional enrichments were considered significant for FDR < 0.01. Functional enrichments in STRING networks, Uniprot keywords, PFAM protein domains, INTERPRO protein domains and features, and SMART protein domains were populated from these comparisons.

## RESULTS

### Generation of the spectral library and DIA assay library

A raw MS2 spectral library was created from the DDA data from 22 samples from both KL and BL populations that were chosen to represent chronic (three weeks) warm (25°C) and cold (7°C) exposure, acute (two hour) warm (28°C) exposure at six and 24 hours post temperature stress and at both elevated (22°C) and control (15°C) recovery temperatures, and acute (2hr) cold stress (4°C) for just the KL population to represent a diverse and representative array of proteins present in stickleback livers after temperature challenge. The assay library includes representatives of both fully plated and low plated morphotypes, as well as representatives from two different types of estuarine habitat, river and lagoon. The DDA search results of these samples were consolidated from different search engines with PEAKS Studio and used for the creation of the MSMS spectral library which initially contained 5,768 proteins, 87,898 peptides, 99,762 precursors and 448,077 transitions.

The spectral library was assembled and then subjected to various quality control measures which greatly reduced the number of transitions, peptides, precursors, and proteins included in the final assay library (Figure 2) as detailed below. The purpose of the filtering steps was to reduce low quality peptides and to remove any ambiguous or redundant peptides that might conflate quantitative peptide analysis. DIA data for all 60 chronic temperature stress samples were used to create the final assay library for this experiment. The number of proteotypic peptides, or peptides unique to a particular protein, are shown in Figure 2e. There were 1714 proteins, 7209 peptides, 8212 precursors, and 40,493 transitions left after all filter steps had been applied. The average number of peptides per protein was 4.2, and the average number of precursors per protein was 4.8. The vast majority (7627) of the precursors in the assay library were represented by five transitions, with 558 precursors represented by four transitions and nine precursors represented by three transitions. 2+ was the most common precursor charge (5244 precursors), then 3+ (2161 precursors), 4+ (399 precursors), and finally 1+ (350 precursors). The five most common transition ions were: y6+ (5044), y7+ (4741), y5+ (4611), y+8 (3984), and y+4 (3765). Any proteins that were significantly different between the fish populations or any treatments were then manually validated, and the absolute final count after all automated filtration and manual validation steps was: 1708 proteins, 7086 peptides, 7928 precursors, and 39,081 transitions. Mass error, retention time reproducibility, and the q-value distributions after the final mProphet model training and peak reintegration serve as quality control validation and are visualized in Figure 3. This assay library can be used in the future to systematically study liver protein abundance for a variety of temperature stress experiments with *G. aculeatus*.

**Figure 2.**
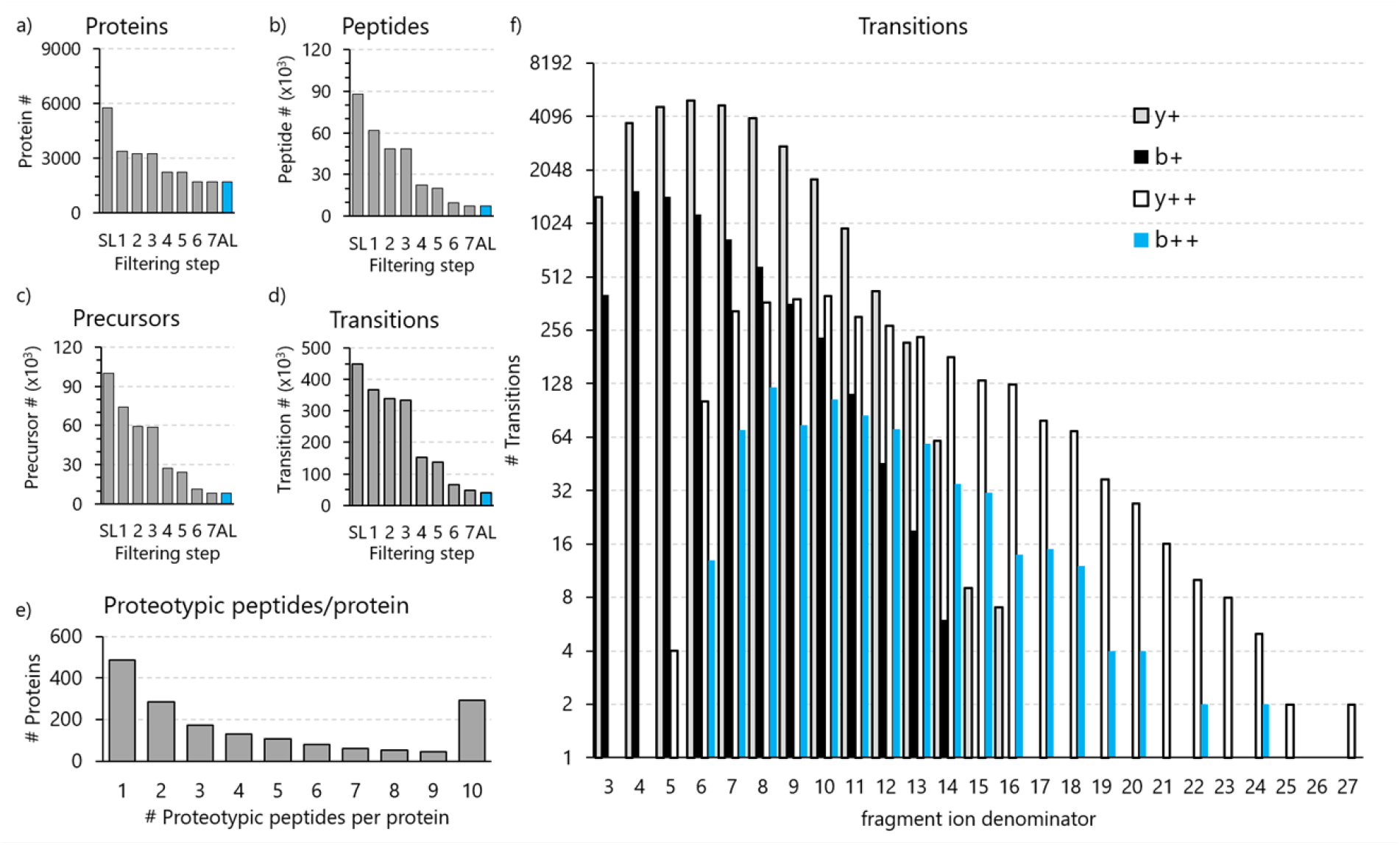
The numbers of a) proteins, b) peptides, c) precursors, and d) transitions are shown throughout the filtration steps used to finalize the liver DIA assay library (AL) from the spectral library (SL). e) Number of unique peptides used for protein quantitation. All proteins were identified by at least two peptides, but for some proteins in the DIA assay library all but one unique peptide was filtered out during the quality control steps. f) Frequency of fragment ion types in the DIA assay library.

**Figure 3.**
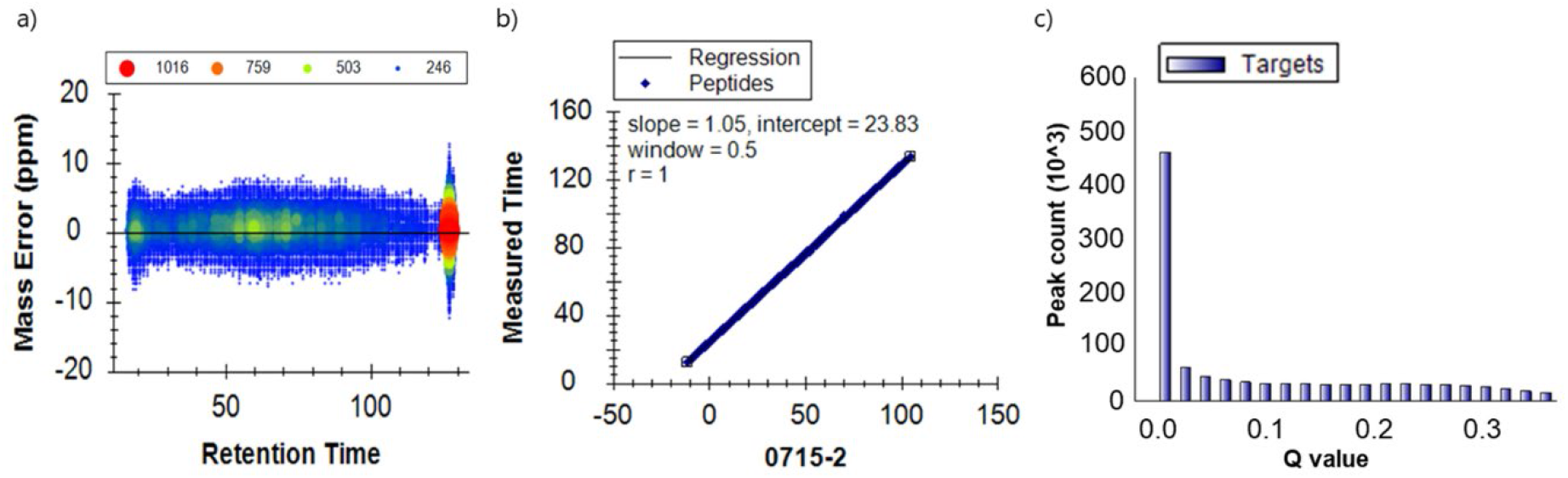
a) Mass error in parts per million (ppm) for all transitions present in the chronic temperature experiment liver assay library as a function of retention time (time of elution from the liquid chromatography column). b) Correlation between measured and predicted retention times. c) Q-values for all peaks of the target peptides in the DIA assay library.

### Population comparisons

#### Overall population comparison (KL vs. BL)

Only a single protein (Figure 4a, Supplemental Table S1; all Supplemental material is available at https://doi.org/10.6084/m9.figshare.14751654) was significantly different between the two populations (all three temperature groups collapsed for each population) and met the fold change threshold (FC > 2). BL had significantly more abundant levels of stromal cell-derived factor 2-like 1 (2.1-fold difference, adjusted p-value = 6.31E-5). There were eight additional proteins significantly higher (adjusted p-value < 0.05) in BL (ribosomal protein L12, heterogeneous nuclear ribonucleoprotein A1a, sorbitol dehydrogenase, thioredoxin, alpha-2-macroglobulin-like 1, heat shock cognate 70, single-stranded DNA binding protein 1, and fibrillarin) and one protein significantly higher in KL (an uncharacterized protein) that did not surpass the minimum fold change cut off (FC > 2) (Supplemental Table S1).

**Figure 4.**
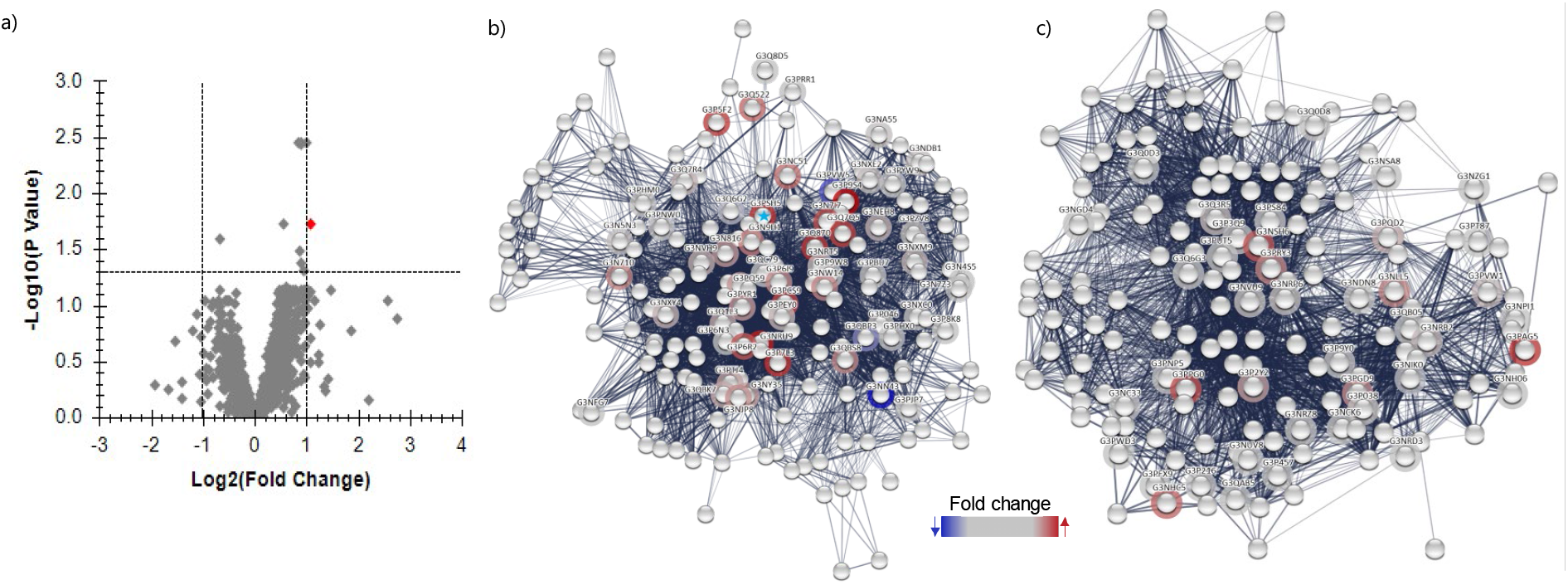
Comparison of liver proteomes in two stickleback populations (KL, N=30, all temperatures collapsed vs. BL, N=30, all temperatures collapsed). a) Volcano plot showing proteins as 1) red diamonds: significantly higher in abundance (FC > 2) and significantly different (adjusted p-value < 0.05), 2) blue diamonds: significantly lower in abundance (FC < 0.5) and significantly different, and 3) grey diamonds: did not meet cut off for both FC and significance requirements. b) & c) Significantly (FDR < 0.01) functionally enriched STRING network clusters, with rings around nodes signifying proteins present in the liver DIA assay library. Rings are colored based on the fold change with red indicating the highest increase in BL relative to KL and blue indicating the highest decrease in BL relative to KL. b) STRING network cluster glycolysis and carbohydrate metabolism (CL:21363, functional enrichment FDR = 2.43E^-7^). The blue star identifies sorbitol dehydrogenase, which was significantly increased in BL over KL at the individual protein level (FC = 1.88, adjusted p-value = 0.0037). c) STRING network cluster AMP-binding, conserved site, and aldehyde dehydrogenase domain (CL:22008, functional enrichment FDR = 3.50E^-4^).

STRING functional enrichment analysis, which was based on fold changes of proteins across the entire liver assay library proteome, identified seven STRING network clusters that were significantly (FDR < 0.01) enriched in BL (Supplemental Table S2). These seven STRING network clusters fell under two main groupings, both of which were elevated in BL (BL > KL), 1) glycolysis and carbohydrate metabolism and 2) AMP-binding and aldehyde dehydrogenase domain. Two representative STRING network clusters are visually depicted in Figure 4b-c, with the corresponding protein list, description, FC, and adjusted-p values in Supplemental Tables S3 and S4. Additional functional enrichments (Supplemental Table S2) that were lower in BL (KL > BL) included the following Uniprot, INTERPRO, PFAM, and SMART protein domains and keywords: winged helix-like DNA-binding domain superfamily, histone H1, and histone H5. Additional Uniprot, INTERPRO, PFAM, and SMART protein domains that were elevated in BL (BL > KL) included oxidoreductase, and NAD(P)-binding domain superfamily.

#### Comparison of populations at 7°C, 15°C, and 25°C

Thirty-five proteins were higher in abundance in BL at 7°C (BL7) and three that were higher in abundance in KL7 passing both the fold change (FC > 2) and significance thresholds (adjusted p < 0.05) (Supplemental Table S1). A volcano plot for the proteins in this comparison are shown in Supplemental Figure 1a and significantly different proteins are visualized in a heat map in Supplemental Figure 2a. Functionally enriched STRING network clusters (Supplemental Table S2) for proteins more abundant in BL7 than KL7 included: low-density lipoprotein (LDL) receptor class A repeat, terpenoid cyclases/protein prenyltransferase alpha-alpha toroid, LDLR class B repeat, lipid transport protein, apolipoprotein A/E, lipoprotein N-terminal Domain, AMP-binding, conserved site, aldehyde dehydrogenase domain, amidohydrolase family, and purine metabolism. Additional functional enrichments for the proteins higher in abundance in BL7 (Supplemental Table S2) included Uniprot keyword oxidoreductase, PFAM, SMART, and INTERPRO domain alpha-2-macroglobulin family, and the additional INTERPRO domain 6-phosphogluconate dehydrogenase.

Proteins higher in abundance in KL7 included COX assembly mitochondrial protein, tubulin-specific chaperone A, and peptidyl-prolyl cis-trans isomerase. No functional enrichments were found from just these three proteins. No significant differences were observed between the two populations at the control temperature of 15°C (KL15 vs. BL15) or after warm acclimation to 25°C (KL25 vs. BL25).

### Effects of temperature on the liver proteome

#### Overall effects of cold acclimation (15°C vs. 7°C)

No proteins were both significantly different and had a fold change greater than two after cold acclimation (15°C vs. 7°C, Figure 5a). L-threonine dehydrogenase was the one protein that was significantly lower at 7°C (adjusted p-value = 0.0004) and alpha-mannosidase was the one protein that was significantly higher at 15°C (adjusted p-value = 0.001), but these proteins did not meet the fold change cut off. There were nine significantly (FDR < 0.01) functionally enriched STRING network clusters (Supplemental Table S2) for 15°C vs. 7°C. STRING networks depleted at 7°C included glycolysis, L-lactate/malate dehydrogenase, tetrahydrofolate dehydrogenase/cyclohydrolase, pyridoxal phosphate-dependent transferase domain 1, carbohydrate metabolism, enolase, and NAD(P)-binding domain. STRING networks enriched at 7°C included ribosome biogenesis, and DEAD/DEAH box helicase. Three of the main STRING network clusters that were significantly functionally enriched, one involving glycolysis and carbohydrate metabolism, a second involving ribosome biogenesis and DEAD/DEAH box helicase, and the last involving pyridoxal phosphate-dependent transferase domain 1 and NAD(P)-binding domain are visually represented in Figure 5b-d. The corresponding protein list, description, FC, and adjusted-p values are provided in Supplemental Tables S5-S7. Additional functional enrichments (15°C > 7°C) included Uniprot keywords glycolysis and pyridoxal phosphate.

**Figure 5.**
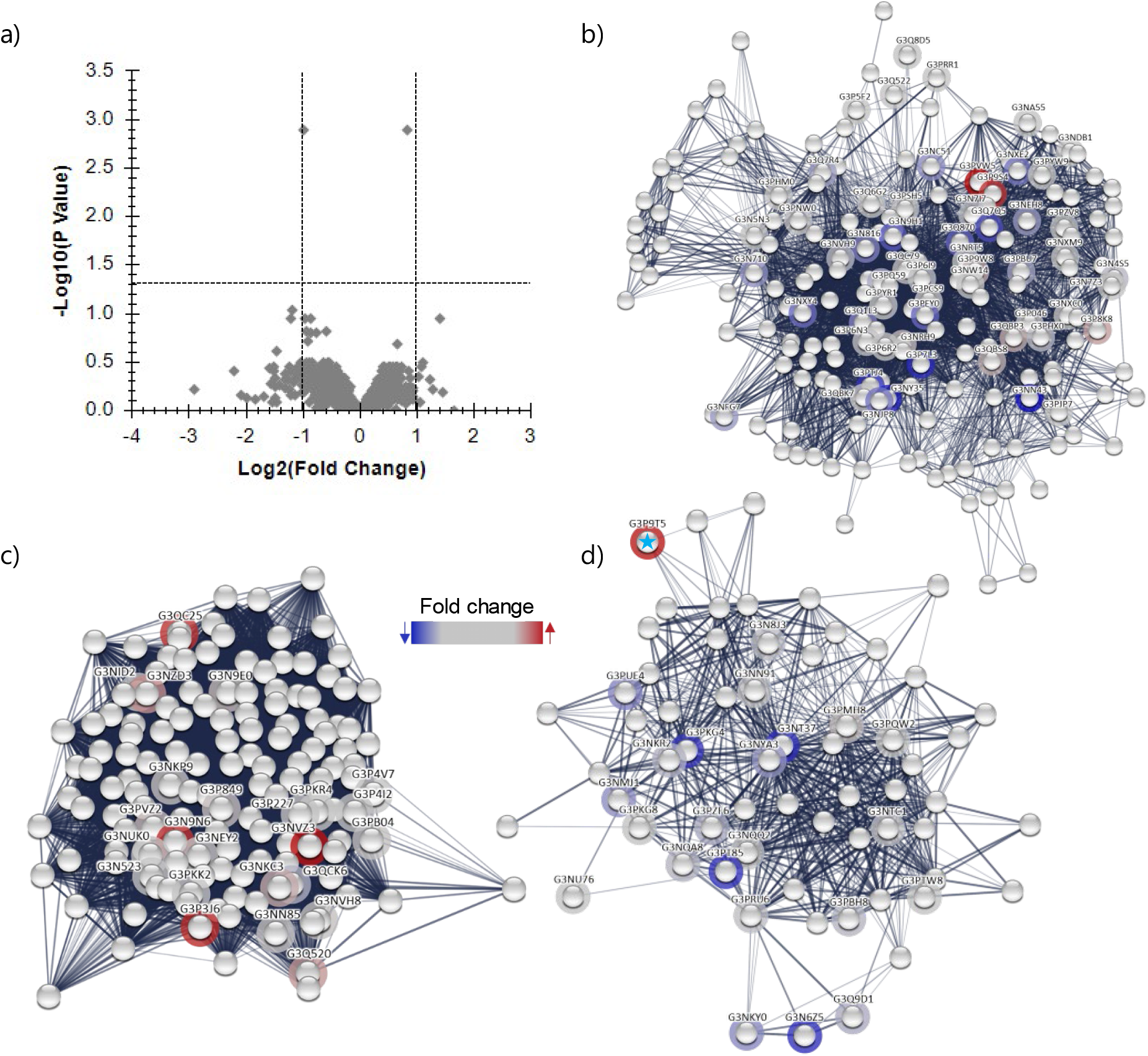
Overall effect of cold stress on the stickleback liver proteome (15°C, N=20, collapsed across both populations vs. 7°C, N=20, collapsed across both populations). a) Volcano plot with proteins depicted as all grey diamonds as none of them met the cut off for both FC (< 0.5 or > 2) and significance requirements (adjusted p-value < 0.05). b)-d) Significantly (FDR < 0.01) functionally enriched STRING network clusters, with rings around nodes signifying proteins present in the liver DIA assay library. Rings are colored based on the fold change relative to all other proteins in the liver set, with red indicating the highest increase in 7°C relative to 15°C and dark blue indicating the maximal decrease in 7°C relative to 15°C. b) Enriched STRING network cluster glycolysis and carbohydrate metabolism (CL:21363, FDR = 0.0039). c) Enriched STRING network ribosome biogenesis and DEAD/DEAH box helicase (CL:16360, FDR = 0.0037). d) Enriched STRING network pyridoxal phosphate-dependent transferase domain 1 and NAD(P)-binding domain (CL:21790, FDR = 0.0054). The blue star identifies sorbitol dehydrogenase (FC = 1.79, adj. p = 0.001).

#### Population-specific effects of cold acclimation (KL15 vs. KL7 and BL15 vs. BL7)

No significant differences were induced by cold acclimation to 7°C in KL fish (KL15 vs. KL7). For BL15 vs. BL7, one protein, alpha-mannosidase, was lower in BL7 and met both significance (adjusted p-value=0.042) and fold change (FC=2.5) requirements (Supplemental Table S1). One additional protein, nucleolar protein interacting with the FHA domain of MKI67 was significantly higher in abundance in BL7 but did not meet the fold change cut off.

#### Overall effects of warm acclimation (15°C vs. 25°C)

A total of 35 proteins were significantly and >2-fold increased at 25°C when combining warm acclimation data for both populations while 51 proteins were significantly and <0.5-fold decreased (Figure 6a-b, Supplemental Table S1). An additional 44 proteins were significantly higher and 108 additional proteins were significantly lower at 25°C, but these proteins did not meet the fold change cut off. Ten functionally enriched STRING network clusters (Supplemental Table S2) grouped into three main categories: 1) core histones H2A/H2B/H3/H4 (elevated at 25°C), 2) ribosomal proteins, including S18, L37, and S30 and translational protein SH3-like (depleted at 25°C), and 3) acyltransferase ChoActase/COT/CPT, and SCP2 sterol-binding domain (depleted at 25°C). Three of the main functionally enriched (FDR < 0.01) STRING network clusters representing each of the three main groupings are visually represented in Figure 6d-f. The corresponding protein list, description, FC, and adjusted-p values are provided in Supplemental Tables S8-S10. Additional functional enrichments at 25°C included Uniprot keywords chromosome and DNA-binding, and PFAM, INTERPRO, and SMART protein domains histone H1 and H5 family, and winged helix-like DNA-binding domain superfamily. Additional functional depletions at 25°C included ribosomal protein S5 domain and thiolase-like.

**Figure 6.**
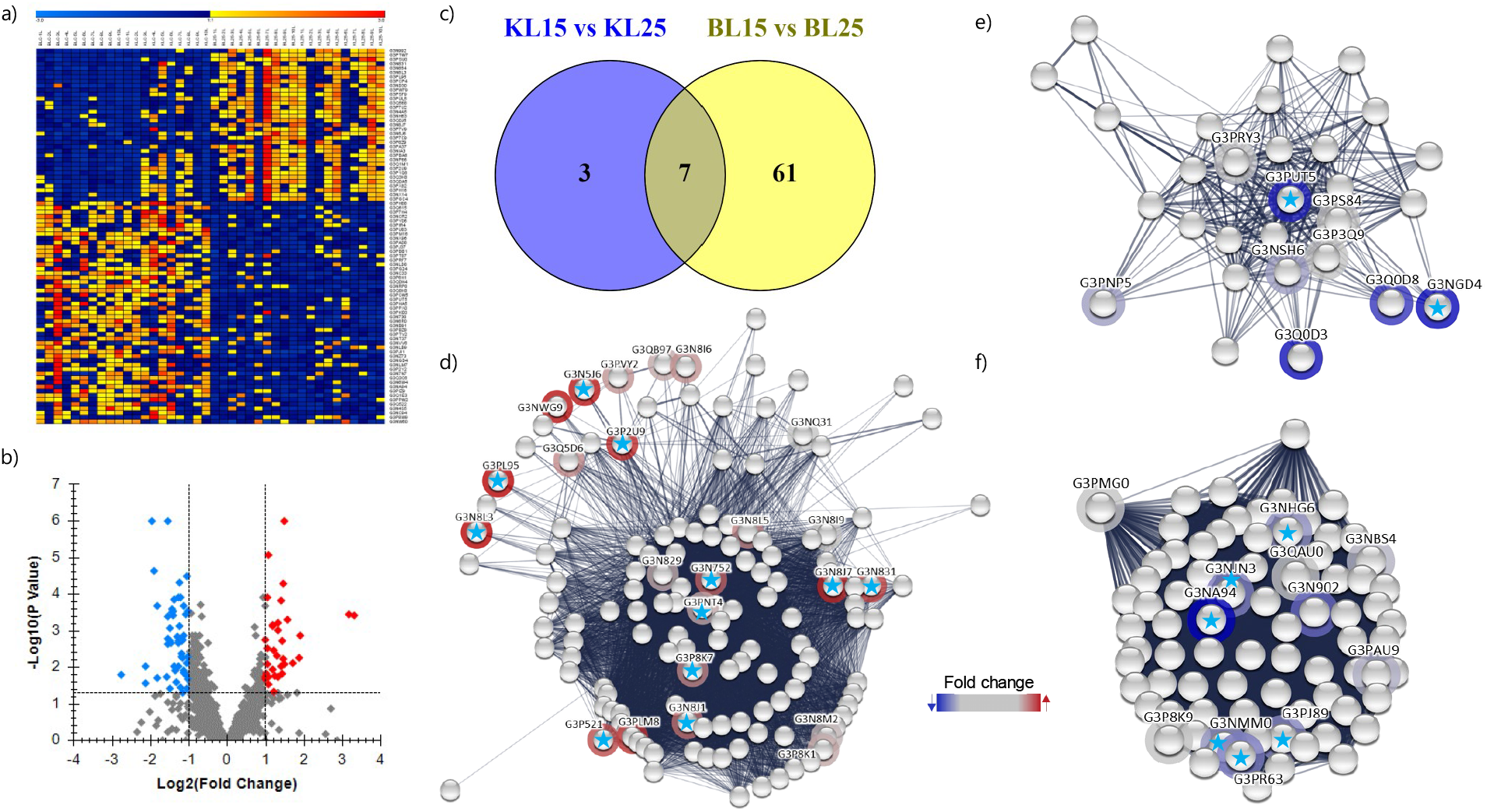
Overall effect of warm stress on the stickleback liver proteome (15°C, N=20, collapsed across both populations vs. 25°C, N=20, collapsed across both populations). a) Heat map depicting significantly (adjusted p-value < 0.05) up and down regulated proteins for all biological replicates. Yellow to red coloring represents proteins with a higher abundance, with red having the highest abundance. Dark blue to light blue represents proteins with a lower abundance, with light blue having the lowest abundance. b) Volcano plot showing proteins depicted as 1) red diamonds: significantly higher in abundance (FC > 2) and significantly different (adjusted p-value < 0.05), 2) blue diamonds: significantly lower in abundance (FC < 0.5) and significantly different, and 3) grey diamonds: did not meet cut off for both FC and significance requirements. c) Venn diagram depicting number of significantly different proteins between the KL15°C vs. KL25°C comparison and the BL15°C vs. BL25°C comparison. d)-f) Significantly (FDR < 0.01) functionally enriched STRING network clusters, with rings around nodes signifying proteins present in the liver DIA assay library. Rings are colored based on the fold change relative to all other proteins in the liver set, with red indicating the highest increase in 25°C relative to 15°C and blue indicating the maximal decrease in 25°C relative to 15°C. Blue stars indicate individual proteins that were significantly different in abundance between 15°C vs. 25°C d) Enriched STRING network cluster, core histone H2A/H2B/H3/H4, and histone H4 (CL:11311, FDR = 8.88E^-9^). e) Enriched STRING network cluster, acyltransferase choactase/COT/CPT, and SCP2 sterol-binding domain (CL:22217, FDR = 0.0027). f) Enriched STRING network cluster, ribosomal protein, and ribosomal protein S18 (CL:16051, FDR = 0.0023).

#### Population-specific effects of warm acclimation in KL fish

Five proteins were significantly and >2-fold elevated in abundance in KL fish acclimated to 25°C (KL25): heterogeneous nuclear ribonucleoprotein D, ATP synthase, H+ transporting, mitochondrial Fo complex, subunit F6, OCIA domain containing 1, and two uncharacterized proteins. Five proteins were also significantly and <0.5-fold reduced in abundance in KL25: Leukocyte cell-derived chemotaxin 2 like, phytanoyl-CoA 2-hydroxylase, peptidylprolyl isomerase, mitochondrial ribosomal protein S16, and ribonuclease T2) (Supplemental Table S1). There was one additional protein significantly higher for KL25 (heat shock protein family [HSP40] member B1b) and lower for KL25 (adenylate kinase 2, mitochondrial) that did not meet the fold change cut off. No functional enrichments were found from these proteins. A volcano plot for the proteins in this comparison is shown in Supplemental Figure S1b and significantly different proteins are visualized in a heat map in Supplemental Figure S2b.

#### Population-specific effects of warm acclimation in BL fish

Seventeen proteins were significantly and >2-fold elevated in BL fish acclimated to 25°C (BL25) while 51 proteins were significantly and <0.5-fold reduced in BL25 (Supplemental Table S1). A volcano plot for the proteins in this comparison is shown in Supplemental Figure S1c and significantly different proteins are visualized in a heat map in Supplemental Figure S2c.

Significant functional enrichments (Supplemental Table S2) in BL25 included STRING network clusters core histone H2A/H2B/H3/H4, Uniprot keyword chromosome, PFAM protein domain linker histone H1 and H5 family, INTERPRO Protein domains and features histone H5 and linker histone H1/H5, domain H15, and SMART protein domains histone families 1 and 5. Significant functional depletions (Supplemental Table S2) in BL25 included seven STRING network clusters pertaining to ribosomal protein and protein biosynthesis, several proteinase inhibitors, peptidase S1A, coagulation factors VII/IX/X/C/Z, cystatin, cathepsin D, fibrinogen, and PAN domain. Additional functional depletions in BL25 included Uniprot keyword RNA-binding and the following PFAM protein domains: RNA recognition motif. (a.k.a. RRM, RBD, or RNP domain), ubiquitin family, various elongation factors, cystatin domain, cyclophilin type peptidyl-prolyl cis-trans isomerase/CLD, and ubiquitin-2 like Rad60 SUMO-like. Finally, 17 INTERPRO protein domains and features were functionally depleted in BL25: RNA recognition motif, nucleotide-binding, RNA-binding, multiple different elongation factors, ubiquitin, cystatin, thiolase-like, cyclophilin-type peptidyl-prolyl cis-trans isomerase, ribosomal protein S5, and transcription factor GTP-binding.

#### Population overlap of warm acclimation effects

Seven proteins overlapped in being significantly higher or lower in abundance after 25°C acclimation in both KL and BL populations (Figure 6c). The increased proteins are 1) heterogeneous nuclear ribonucleoprotein, 2) ATP synthase, H+ transporting, mitochondrial Fo complex, subunit F6, and 3) OCIA domain containing 1. The decreased proteins are: 1) leukocyte cell-derived chemotaxin 2 like, 2) phytanoyl-CoA 2-hydroxylase, 3) peptidylprolyl isomerase, and 4) ribonuclease T2. No significant functional enrichments were identified from these seven proteins.

#### Overall differences between cold and warm acclimation (7°C vs. 25°C)

A total of 77 proteins were significantly and >2-fold higher in abundance at 25°C vs. 7°C and 72 proteins were significantly and <0.5-fold lower in abundance at 25°C vs. 7°C (Figure 7, Supplemental Table S1). Additionally, there were 73 more proteins that were significantly increased at 25°C and 79 more proteins that were significantly increased at 7°C, but did not meet the fold change cutoff. Results are similar to the comparison of the cold and warm comparisons to the control temperature (15°C), so networks are not depicted. Functionally enriched STRING network clusters (Supplemental Table S2) included five pertaining to core histones H2A/H2B/H3/H4 (25°C > 7°C), and one network cluster for ribosomal protein and ribosomal protein L37/S30 (7°C > 25°C). UniProt keywords that were functionally enriched at 25°C included chromosome, DNA-binding, and nucleosome core, while RNA-binding was depleted at 25°C. PFAM, INTERPRO, and SMART domains that were functionally enriched at 25°C included linker histone H1 and H5 family, histone fold, and histones H2A/H2B/H3 while RNA binding domain, RNA recognition motif, elongation factor G, nucleotide-binding alpha-beta plait domain superfamily, thiolase-like, and ribosomal protein S5 were depleted at 25°C.

**Figure 7.**
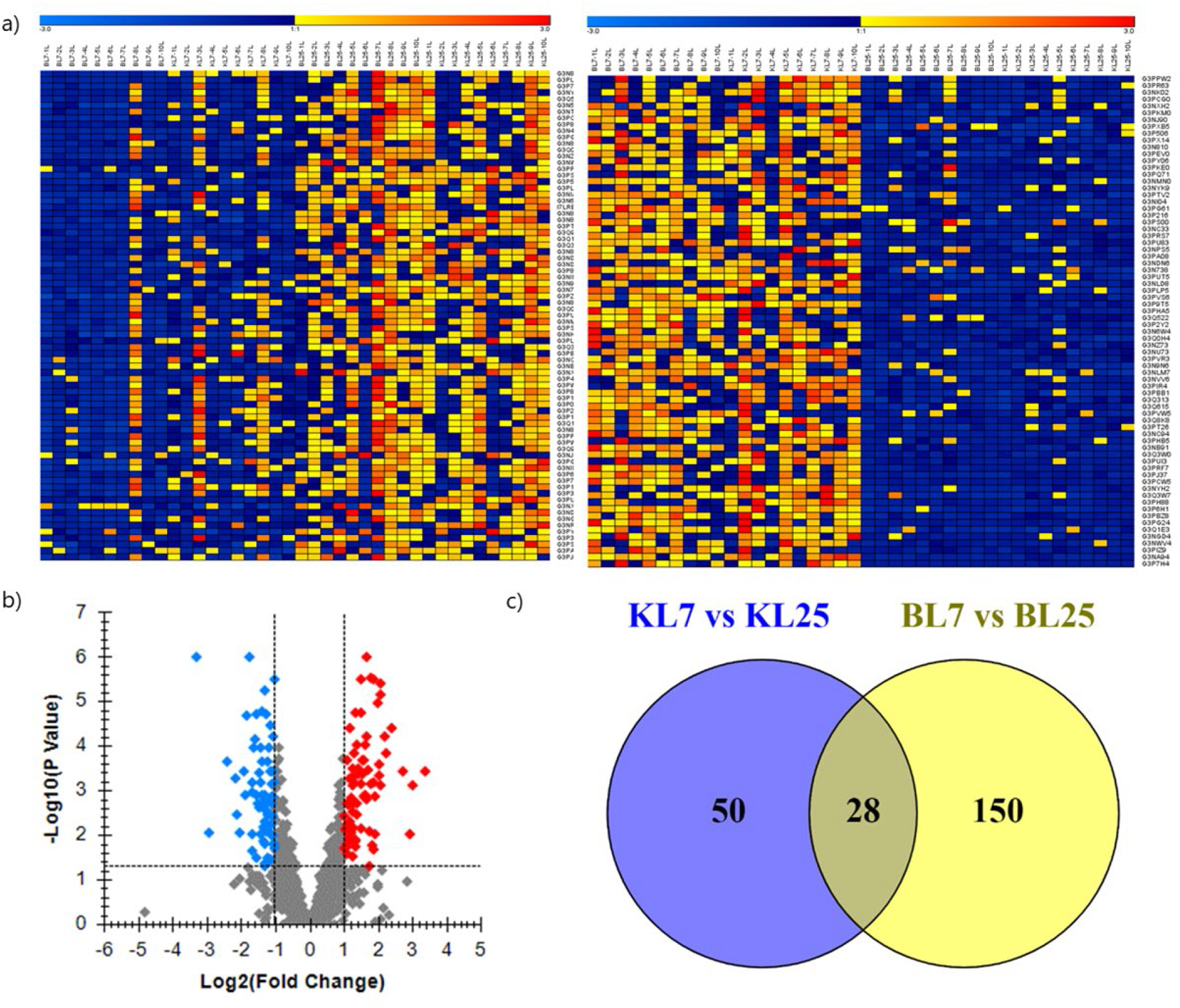
Comparison of cold versus warm liver proteomes in two stickleback populations (7°C, N=20, collapsed across both populations vs. 25°C, N=20, collapsed across both populations). a) Heat map depicting significantly (adjusted p-value < 0.05) up and down regulated proteins for all biological replicates. Yellow to red coloring represents proteins with a higher abundance, with red having the highest abundance. Dark blue to light blue represents proteins with a lower abundance, with light blue having the lowest abundance. b) Volcano plot showing proteins depicted as 1) red diamonds: significantly higher in abundance (FC > 2) and significantly different (adjusted p-value < 0.05), 2) blue diamonds: significantly lower in abundance (FC < 0.5) and significantly different, and 3) grey diamonds: did not meet cut off for both FC and significance requirements. c) Venn diagram depicting number of significantly different proteins between the KL7°C vs. KL25°C comparison and the BL7°C vs. BL25°C comparison.

#### Population-specific differences between cold and warm acclimation in KL fish

Forty-nine proteins were significantly and >2-fold elevated in KL fish at 25°C (KL25) and 29 proteins were significantly and >2-fold higher in abundance in KL7 (Supplemental Table S1). A volcano plot for the proteins in this comparison is shown in Supplemental Figure S1d and significantly different proteins are visualized in a heat map in Supplemental Figure S2d. Functional enrichments for the 49 proteins higher in abundance in KL25 included the following STRING network clusters (Supplemental Table S2): glycolysis, enolase, phosphoglycerate mutase 1, carbohydrate metabolism, calponin repeat, caldesmon, annexin A2 and A11, FAD dependent oxidoreductase, D-isomer specific 2-hydroxyacid dehydrogenase, catalytic domain, and core histones H2A/H2B/H3/H4. Functionally enriched UniProt keywords elevated in KL25 included chromosome and nucleosome core. Additional functional enrichments (KL25 > KL7) for PFAM, INTERPRO, and SMART protein domains included core histones H2A/H2B/H3/H4, enolase, TIM barrel domain, and tropomyosin.

Functional enrichments (Supplemental Table S2) for the 29 proteins significantly higher in abundance in KL7 included STRING network clusters of ribosomal proteins, RNA recognition motif domains and mRNA processing, translation protein SH3-like domain superfamily, and protein biosynthesis. Additional functional domains and keywords that were depleted in KL7 (KL25 > KL7) pertained to RNA-binding, RNA recognition motif nucleotide-binding alpha-beta plait domain superfamily.

#### Population-specific differences between cold and warm acclimation in BL fish

Enforcing a fold change cut off of FC > 2, eighty-two proteins were significantly increased in BL fish acclimated to 25°C (BL25) while 96 proteins were significantly higher in abundance in BL7 (Supplemental Table S1). A volcano plot for the proteins in this comparison is shown in Supplemental Figure S1e and significantly different proteins are visualized in a heat map in Supplemental Figure S2e. Functional enrichments (Supplemental Table S2) for the 82 proteins increased in BL25 included STRING network clusters pertaining to core histones H2A/H2B/H3/H4 and H5, HMG box A DNA-binding domain, calponin repeat, and caldesmon. Functionally enriched Uniprot keywords in BL25 included chromosome, DNA-binding, nucleosome core, and the nucleus. Functionally enriched PFAM, INTERPRO, and SMART protein domains in BL25 involved linker histone H1 and H5 histone H2A, Histone H5, and winged helix-like DNA-binding domain superfamily.

Functional depletions (Supplemental Table S2) for the 96 proteins in BL7 vs. BL25 included the following STRING network clusters: ribosomal protein, protein biosynthesis, heterogeneous nuclear ribonucleoprotein C, HnRNP-L/PTB, HSP70 protein, DnaJ C terminal domain, hnRNP A0, RNA recognition motif translation protein SH3-like domain superfamily, fatty acid hydroxylase, sterol biosynthesis, and mRNA processing. Uniprot Keywords that were depleted in BL25 included RNA-binding, cytoplasm, cytoskeleton, nucleotide-binding, protein biosynthesis, sterol biosynthesis, microtubule, lipid metabolism, lipid biosynthesis, ATP-binding, and tricarboxylic acid cycle. Moreover, numerous elongation factors, RNA recognition motif, tubulin/FtsZ family, GTPase, nucleotide-binding alpha-beta plait domain superfamily, RNA-binding domain superfamily, tubulin, ribosomal proteins, transcription factor, translation protein beta-barrel domain superfamily, K homology domain, and thiolase-like proteins were functionally depleted in BL25 vs. BL7.

#### Population overlap of cold- versus warm-acclimation effects

Twenty-eight proteins that were significantly regulated in the same direction during warm acclimation of both populations (17 increased, 11 decreased) (Figure 7c). Functional enrichments (Supplemental Table S2) for these 28 overlapping significant proteins included calponin repeat, and caldesmon, annexin A2, and annexin A11, and core histone H2A/H2B/H3/H4, and histone H4. Additional function enrichments included Uniprot keywords chromosome, PFAM protein domains C-terminus of histone H2A, INTERPRO protein domains histone H2A, C-terminal domain, and histone H2A conserved site, and SMART protein domains histone 2A and calponin homology domain.

## DISCUSSION

### Proteins involved in protein homeostasis fuel higher metabolic need in BL sticklebacks

As a population, BL had a higher number of elevated proteins over KL and had more significantly different proteins for each within population temperature comparison than KL. Although a small number of significantly regulated proteins were shared between both populations, our study revealed clear differences in how the two populations handled chronic temperature stress. The network analyses helped elucidate some of these differences. Chronic cold temperature challenge yielded most differences between the two populations.

Regardless of temperature, all but one of the proteins that were significantly different between the populations were higher in abundance in the BL population. Many of these proteins are either directly or peripherally involved in proteostasis. Stromal cell-derived factor 2-like 1 (SDF2L1) is localized in the endoplasmic reticulum (ER) and, in mice, it has been suggested to increase the amount of time available for misfolded proteins to regain their correct conformation and that it acts as a regulator in the ER stress response (37, 38). The constitutively expressed molecular chaperone heat shock cognate 70 (hsc70) is involved in a range of protein homeostasis functions such as *de novo* protein folding, protein translocation, protein assembly and disassembly, regulation of protein activity, protection from proteolysis, and coordinating with other smaller molecular chaperones (39, 40). Thioredoxin is a ubiquitous antioxidant found in all cell types and organisms (41). Alpha-2-macroglobulin-like 1 is involved in the immune response through the complement and coagulation cascades and was elevated in grass carp (*Ctenopharyngodon idellus*) after 48 hour exposure to high temperature (42). 60S ribosomal protein L12 is at the core of translation and catalyzes the formation of peptide bonds (43) while heterogeneous nuclear ribonucleoprotein A1a is involved in mRNA splicing and stability as well as overall regulation of translation (44). Fibrillarin is a component of a nucleolar ribonucleoprotein involved in rRNA processing (45), and single-stranded DNA binding protein 1 is integral for processes such as DNA replication and repair (46). Overall, these proteins elevated by temperature in BL represent aids for protein folding or refolding, reactive oxygen species (ROS) scavenging, immune response, transcription, and translation. Increases in these processes signify a higher metabolic demand. In support of this notion, sorbitol dehydrogenase was significantly higher in abundance in BL over KL and from the network analysis, glycolysis and carbohydrate metabolism were functionally enriched with proteins aligning to this network being elevated in BL over KL.

The main difference between the two populations was revealed by cold acclimation. Again, BL fish contributed the vast majority of significantly elevated proteins, and conversely, KL fish had a significantly lower abundance of many proteins. Molecular chaperones heat shock cognate 70 and heat shock 60 protein 1 were elevated in BL7 as were the previously discussed proteins stromal cell-derived factor 2-like 1 and single-stranded DNA protein 1. Two aldehyde dehydrogenases (ALDH) from family 8 and 2 were also higher in BL7 and ALDH was found to be functionally enriched as well. These proteins are oxidative stress proteins and aldehyde dehydrogenase was similarly elevated after cold acclimation in the mussel *Mytilus trossulus* (47). Aspartate aminotransferase (ALT) and alanine-glyoxylate aminotransferase 2 were both more abundant in BL7 and play a role in glutamate metabolism (48). Both of these proteins also serve as biomarkers for liver damage (49), as can alpha-2-macroglobulin, which was both functionally enriched and significantly elevated at the individual protein level and may explain the increase in the ROS-scavenging ALDH proteins. Both alpha and beta chain tubulin proteins and one tubulin-specific chaperone were significantly elevated in BL7 over KL7. Cold acclimated gilthead seabream (*Sparus aurata*) also had elevated levels of tubulin, which comprise the microtubule network forming the cytoskeleton and are thought to play a role in the cellular stress response, although the exact mechanism is unknown (50, 51). Additionally, in a study examining two mussel congeners, the warm-adapted congener had increased abundances of tubulin after cold acclimation (47). Glycerol-3-phosphate dehydrogenase (GPDH) was one of the proteins significantly elevated in BL7 over KL7. GPDH activity increased at low temperatures in rainbow smelt (*Osmerus mordax*) (52, 53) and snow trout (*Schizothorax richardsonii and Schizothorax niger*) (54) and resulted in an increase in glycerol, which can act as an antifreeze. From our functional enrichment analysis, 6-phosphogluconate dehydrogenase (6PGD) was identified twice from the INTERPRO database. 6PGD is a part of the pentose phosphate pathway (PPP) and produces NADPH which can reduce ROS via the glutathione system, so 6PGD can be considered an antioxidant enzyme (55). Consistent with a role of the PPP, transaldolase is another protein elevated by cold acclimation in our study (56).

In zebrafish (*Danio rerio*), cold resistance is conferred through lipid catabolism and autophagy (57). In the BL7 group, autophagy-related protein 3 was in higher abundance along with various proteins involved with lipid metabolism, such as GPDH, 3-hydroxyisobutyrate dehydrogenase, and lipid transport (apolipoprotein Ea). The autophagy-related protein 3 and lysosomal enzyme N-acetylglucosamine-6-sulfatase suggest the increased need in the BL7 group to degrade damaged molecules, which is consistent with an increase in ROS related damage. Aconitate hydratase catalyzes the isomerization of citrate to isocitrate in the citric acid cycle and is sensitive to ROS (58). There are clear differences between the two populations, with BL exhibiting higher abundances of proteins involved in carbohydrate, amino acid, and lipid metabolism, as well as an increase in antioxidant proteins and molecular chaperones. These observations are consistent with the conclusion that higher metabolic demand triggers increased ROS production during cold acclimation of BL fish.

Overall, there could be several reasons for these population differences. The BL population might have a more uniform response among individuals in the population thus leading to more significant differences, while the KL population may have greater within-population variation in how individuals deal with temperature stress. Alternatively, BL might have more energy available to devote to processes that demand higher metabolic rates or simply be able to alter metabolic flux more easily. A third possibility is that the KL population was able to achieve a state of homeostasis faster than the BL population and that after three weeks, concentrations of many proteins had returned to baseline levels in KL while BL was still mounting a response to the chronic stress. KL and BL might also be employing different strategies, one that requires more upkeep and energy versus one that is less responsive but has a lower energetic cost.

### Both populations re-establish liver proteome homeostasis during chronic cold acclimation

Despite differences at 7°C between the two populations, there were no significant effects of cold acclimation on individual proteins irrespective of whether the populations were combined or analyzed within population. The lower limit for marine populations is around 4°C and for freshwater populations it is around 1°C (59). However, even in freshwater lakes farther north in British Columbia, 4°C is the mean monthly temperature for winter months (59). Functional enrichments pertaining to glycolysis, ribosome biogenesis, and RNA metabolism (DEAD/DEAH box helicase) hint at some differences between 15°C and 7°C, but for the most part, it appears that the populations were able to re-establish homeostasis after the initial temperature shock at 7°C. This is likely a temperature that both populations would experience in the wild. Given that water has the highest density at 4°C, it would be unrealistic to expose these populations to conditions at much lower than 4°C. Whether a cold-acclimation to 4°C results in a more pronounced response than cold-acclimation to 7°C remains to be explored in future studies.

### Chronic warm acclimation increases chromatin regulation and transcriptional control while reducing translation and fatty acid metabolism proteins

Comparing fish acclimated to 25°C versus 7°C yielded similar results as those discussed for 7°C vs. 15°C above and for 25°C vs. 15°C below, but with a much higher statistical significance. Most proteins that are significantly increased at 25°C compared to 15°C controls are histones and proteins that regulate transcription or chromatin structure. Host cell factor C1a has been linked to cell proliferation, gluconeogenesis promotion, and regulation of transcription (60). Two ribonucleoproteins were higher in abundance, including SAP domain containing ribonucleoprotein and heterogeneous nuclear ribonucleoprotein D, which was elevated at 25°C for the overall temperature and population-specific comparisons and serves as a transcriptional repressor (61). Serpine1 mRNA binding protein 1b is involved in mRNA stability (62, 63). Ataxin 2-like protein is a component of stress granules in mammals, but is evolutionarily conserved across eukaryotes (64), and responds to a variety of stressors via mRNA regulation with links to mRNA degradation (65). One study on carp (*Cyprinus carpio*) found that the expression of four H2A variants were enriched during the summer, and that a ubiquitylated variant regulated chromatin structure (66). Another study also demonstrated large changes in nucleolar structure and the expression of ribosomal genes with acclimatization to seasonal changes in carp (*C. carpio*) liver (67). Besides numerous histone proteins with significantly higher abundance at 25°C, functional enrichment of networks involving core histones (including 11 proteins that were significantly more abundant in 25°C) as well as functional enrichment of chromosome and DNA binding point to a key role of chromatin regulation for warm acclimation. It is likely that there is epigenetic regulation occurring in various histones to both alter the chromatin structure and regulate transcription and translation to acclimate to the increased temperature. Future studies may examine post-translational modifications (PTMs) associated with these elevated proteins. PTMs are easily identified in mass spectrometry-based proteomics but quantifying them is not trivial and exceeded the scope of this study.

Additional proteins elevated at 25°C included paxillin, which is a molecular scaffold or adaptor protein that regulates cellular responses to changes in the environment (68). Paxillin may interact with other structural proteins that were elevated at 25°C, such as tropomyosin 1 and calponin, to increase cytoskeletal stability. Peptide-methionine (R)-S-oxide reductase can be considered an antioxidant enzyme that converts methionine sulfoxide back to the amino acid methionine, one of the easiest amino acids to oxidize (69). Two isoforms of Tpd52 like protein 2b were also elevated at 25°C. Tpd52 associates with lipid droplets and likely signifies changes in lipid storage (70).

Functional enrichment analysis for proteins that were decreased at 25°C, revealed ribosomal and thiolase like proteins, suggesting a decrease in translation (including significantly reduced mitochondrial ribosomal proteins S16, L41, L24, L14, S18A, and L27; Supplemental Table S10) and alteration in fatty acid metabolism (including significantly lower abundance proteins: fatty acid synthase, acetyl-CoA acetyltransferase 2, and sterol carrier protein 2a). Phytanoyl-CoA 2-hydroxylase, another protein involved in fatty acid metabolism, is involved in the oxidation of 3-methyl branched fatty acids (71). There were also three elongation factors (eukaryotic translation elongation factor 2b, elongation factor like GTPase1, and eukaryotic translation elongation factor 2a) and two splicing factors (serine/arginine-rich splicing factor 4 and serine/arginine-rich splicing factor 2b) that were less abundant at 25°C, adding more evidence of a decrease in translation.

Leukocyte cell-derived chemotaxin 2-like protein was lower in abundance in the overall comparison and for both population comparisons. Leukocyte cell-derived chemotaxin (LECT2) modulates immune function and inflammatory pathways and serum levels are indicative of liver fat content (72). Chronic stress can divert energy away from the immune system (30), and the lower levels of this protein could also indicate a depletion of liver fat content that would likewise explain a decrease in various proteins linked to fatty acid metabolism.

There were many more significantly different proteins in the population-specific comparisons that support the above conclusions regarding the overall warm-acclimation effect. Proteins that were increased in the BL population clustered into the same functional categories as those discussed above for overall differences, i.e. histone and chromatin structure, and transcription. Proteins that were significantly elevated at 25°C only in the BL population (BL25 > BL15) include centromere protein V, SUB 1 homolog, transcriptional regulator 1, and pleckstrin homology domain containing, family A member 6 (PLEKHA6). The overexpression of centromere protein V can lead to hypercondensation of certain types of heterochromatin (73). SUB 1 homolog, transcriptional regulator b is inducible by oxidative stress and protects DNA from oxidative damage when it is exposed or partially unwound (74). Therefore, these proteins support our conclusion of significant effects of temperature acclimation on chromatin structure and transcriptional regulation.

Functions of the proteins that were significantly lower in abundance only in BL fish acclimated to 25°C (BL15 > BL25) also reflected functional categories that were depleted at 25°C overall in both populations. Functional enrichment analysis revealed significant depletion of ribosomal proteins in cumulative BL25+KL25 and BL25 only datasets. Beyond just ribosomal proteins, there were also individual proteins with significantly lower abundance at BL25 that are involved more generally in protein biosynthesis (eukaryotic translation elongation factor 2b, RNA binding protein S1 serine-rich domain, nascent polypeptide associated complex subunit alpha, ribosomal protein L37, eukaryotic translation initiation factor 3 subunit G, and eukaryotic translation elongation factor 1 beta 2). Two peptidylprolyl isomerase proteins were also lower in abundance. These proteins help with folding newly synthesized proteins, but also play a role in the immune system, cell cycle control, and transcriptional regulation (75). Two lysosomal proteins (legumain and acid phosphatase 2, lysosomal) were likewise lower in abundance in BL25. Filamin B is an actin-binding cytoskeletal protein, but also binds RNA and decreased transcript levels of filamin B have resulted in downregulation of apoptosis and genes that function in immunity and inflammation (76), which is consistent with some of the functions discussed above. Besides the common proteins involved in lipid metabolism that were significantly lower in abundance in both populations as mentioned above (sterol carrier protein 2a, phytanoyl-CoA 2-hydroxylase, fatty acid synthase), an additional lipid metabolism protein (lipase) was significantly reduced in the BL population. Lipase is involved in triglyceride regulation (77). Another protein of note that was only significantly reduced at 25°C in BL fish (BL15 > BL25) was thioredoxin domain containing 17, which is involved in cellular redox homeostasis (78).

### Proteome signatures provide comprehensive insight into mechanisms of environmental acclimation

While individual proteins can serve as good bioindicators for injury, disease, or other conditions, proteomic signatures, which encompass the patterns of multiple proteins of interest, have the potential to give specific insight into the physiological state or condition of an organism (79). With enough data sets, the repeated co-expression patterns of a wide range of proteins could potentially give more context and specificity to the type and degree of a particular stressor.

While our data is derived from only two populations from one species, it provides a very detailed snapshot of the molecular phenotypes of these organisms and can identify the strategies and mechanism utilized to overcome a change in the environment. Proteins often have numerous roles, so connecting overall changes in protein abundance helps disentangle the specific pathways that are activated or suppressed and to identify new networks and connections.

## Author contributions

BL designed, performed, and analyzed the experiments, and wrote the paper. DK contributed to experimental design and proteomics data acquisition and analysis, and revised the paper.

## DATA ACCESSIBILITY

All proteomics data and metadata are accessible at the following repositories: MassIVE (ID=MSV000087672, ProteomeXchange ID=PXD026823) for all DDA data and Panorama Public (https://panoramaweb.org/bbl02.url, ProteomeXchange ID=PXD024617) for the DIA assay library and all DIA data.

